# Targeted H3K9 acetylation at lncSox1 promoter by cell type-specific epigenome editing promotes intermediate progenitor proliferation in developing mouse cortex

**DOI:** 10.1101/2024.11.05.621758

**Authors:** Godwin Sokpor, Pauline Antonie Ulmke, Hoang Duy Nguyen, Linh Pham, Princy Kakani, Van Trung Chu, Sebastian J. Arnold, Beate Brand-Saberi, Huu Phuc Nguyen, Tran Tuoc

## Abstract

The distribution and level of epigenetic (chromatin) marks have implications for differential regulatory effects at specific gene loci. Herein, we applied a protocol which combines *in vivo* electroporation and a CRISPR-dead (d)Cas9 system to probe and edit a specific chromatin mark in the epigenome of intermediate progenitor cells (IPCs) in developing mouse cortex. We found that the promoter of *lncSox1*, a long non-coding gene, is a key genomic locus for H3K9 acetylation (H3K9ac) during IPC amplification. CRISPR-dCas9–mediated addition of H3K9ac at *lncSox1* promoter resulted in lncSox1 upregulation, with attendant increase in IPC pool and augmented neurogenesis. Thus, we have identified dynamic regulation of lncSox1 as a major downstream target of H3 acetylation, and as part of an epigenetic mechanism involved in IPC proliferation during neocortex expansion. This finding is a proof-of-concept that our epigenome editing-based method can be used for manipulating specific epigenetic effectors to determine their (neuro)biological significance.

**MOTIVATION:** The abundance of basal progenitors is critical for cortical neurogenesis during brain development. There is growing interest in identifying how specific epigenetic factors regulate the genesis and expansion of basal progenitor cell sub-populations, including intermediate progenitor cells. We established a protocol that allowed us to identify the involvement of a long non-coding RNA (*lncSox1*) in regulating the proliferation of intermediate progenitor cells under the influence of H3K9 acetylation (H3K9ac). By enhancing H3K9ac at the promoter region of *lncSox1* using a CRISPR-dCas9–mediated gene-editing tool, we were able to determine that lncSox1 upregulation is a downstream effect of H3K9ac acetylation and is necessary for intermediate progenitor pool amplification during cortical development.

**Highlights:**

– Identification of H3 acetylation-dependent expression of ncRNAs in developing cortex.
– Establishment of cell Cre/LoxP and CRISPR-dCas9-dependent H3K9ac epigenome editing.
– CRISPR-dCas9–mediated addition of H3K9ac at lncSox1 promoter resulted in lncSox1 upregulation.
– H3K9 acetylation at lncSox1 promoter enhances proliferation of TBR2-expressing IPCs.
– Targeted epigenome editing revealed lncSox1 as a key regulator of cortical development.

## INTRODUCTION

The epigenome presents an additional layer of gene regulation without genomic sequence alteration, and it is capable of influencing cellular phenotypes in a stable and inheritable manner (Fitz-James and Cavalli, 2022). A prominent feature in the epigenome is the activity of histone modifying-enzymes which can dynamically remodel the epigenetic landscape in response to inherent cellular demands and/or environmental stimuli. Such chromatin modifying factors have enzymatic functions with which they can either install or erase chromatin-associated elements to regulate chromatin accessibility, and therefore modulate gene expression (Li et al., 2018).

One strategy for exploring the biological significance of the dynamic installation of chromatin marks in the epigenome is to rationally control their distribution and study the resulting effect. To that end, many techniques have been developed to manipulate chromatin-related factors, including traditional knockdown, knockout, and overexpression methods. However, these approaches have limitations in terms of efficiency, specificity, and activity. The growing effort to improve such experimental strategies has led to the development of epigenome editing platforms with improved targeting, specificity, and increased efficiency in manipulating single or multiple chromatin marks without disturbing the nucleotide sequences in the genome. In general, epigenome editing tools employ programable DNA-targeting modules which can bring coupled effectors (epiEffectors) to gene loci to carry out desired epigenetic modifications (Kungulovski and Jeltsch, 2016).

In this proof-of-principle study, we applied the CRISPR-Cas9 (Clustered regularly interspaced short palindromic repeats [CRISPR], and CRISPR-associated sequences [Cas9]) platform to effect epigenome editing of histone acetylation at a target gene locus. To conduct cell type-specific epigenome editing, we referred to our recent report that H3 acetylation in mammalian basal progenitors is a notable mechanism underlining the evolutionary complexification of the neocortex (Kerimoglu et al., 2021). Thus, we were guided by this finding to perform experimental manipulation of H3 acetylation in the acetylome of intermediate progenitor cells (IPCs) in the developing mouse cortex using the epigenome editing method.

The acetylation (addition of acetyl group) or deacetylation (removal of acetyl group) of histones by the enzymes acetyltransferases and deacetylases, respectively, are well-known chromatin modifying mechanisms which sculpt the distribution of acetyl histone or chromatin marks along gene bodies (Allfrey et al., 1964). The localization of acetyl chromatin marks, especially at regulatory regions of genes (promoter, enhancers), can modulate gene expression to elicit pertinent cellular phenotypes (Gräff and Tsai, 2013).

We focused on the candidate gene lncRNA Sox1 *(lncSox1)* for our epigenome editing method because it was highlighted to be differentially expressed in TBR2- and TBR2+ cells following brain-wide treatment with the histone deacetylation chemical inhibitor Trichostatin A (TSA) to effect global H3 acetylation upregulation. To test if locally increasing histone acetylation in the promoter region of *lncSox1* could augment its expression, we designed an epigenome editing construct by including a Sox1 small-guidance (sg) RNA (sglncSox1), and a histone acetyltransferase KAT2A module which specifically targets histone 3 (H3) at the lysine position 9 (K9) for acetylation (i.e., H3K9ac). Upon co-transfection with a Cre recombinase encoding plasmid, the designed gLncSox1-dCas9-KAT2A-T2A-eGFP DNA fragment is fully transcribed leading to the assembly of a functional H3K9ac epigenome editing complex. With the resultant epigenome editing platform, we were able to efficiently add the chromatin mark H3K9ac to the promoter region of *lncSox1*, which resulted in increased expression of lncSox1 in cultured Neuro2A cells. *In utero* electroporation (IUE)-mediated transfection of IPCs in the developing mouse cortex with our H3K9ac-enriching *lncSox1* construct resulted in increased IPC proliferation and concomitant increase in neurogenesis. Thus, we present this CRISPR-dCas9-based method as an effective means of editing the epigenome at a targeted gene locus in a specific population of cells to study the associated phenotype.

## RESULTS

### 1. Histone deacetylase inhibition altered the expression of non-coding (nc) RNAs in intermediate progenitors in developing mouse cortex

Neocortex expansion is underscored by the enhanced proliferative potential of intermediate progenitors (IPCs) partly through a likely evolution-driven increase in H3K9 acetylation (H3K9ac) in mammalian IPCs (Kerimoglu et al., 2021). To determine how H3 acetylation affects IPC biogenesis and/or proliferative capacity, we considered analyzing the IPC epigenome and transcriptome for downstream factors attributable to the prominence of IPCs in brain evolution. To achieve this, we globally increased H3 acetylation in the embryonic mouse brain by treatment with the selective class I/II histone deacetylase inhibitor (HDACi) Trichostatin A (TSA) via a daily intraperitoneal injection of pregnant wild-type mouse from E12.5 to E16.5 (Fig. 1A) (Kerimoglu et al., 2021).

**Figure 1.**
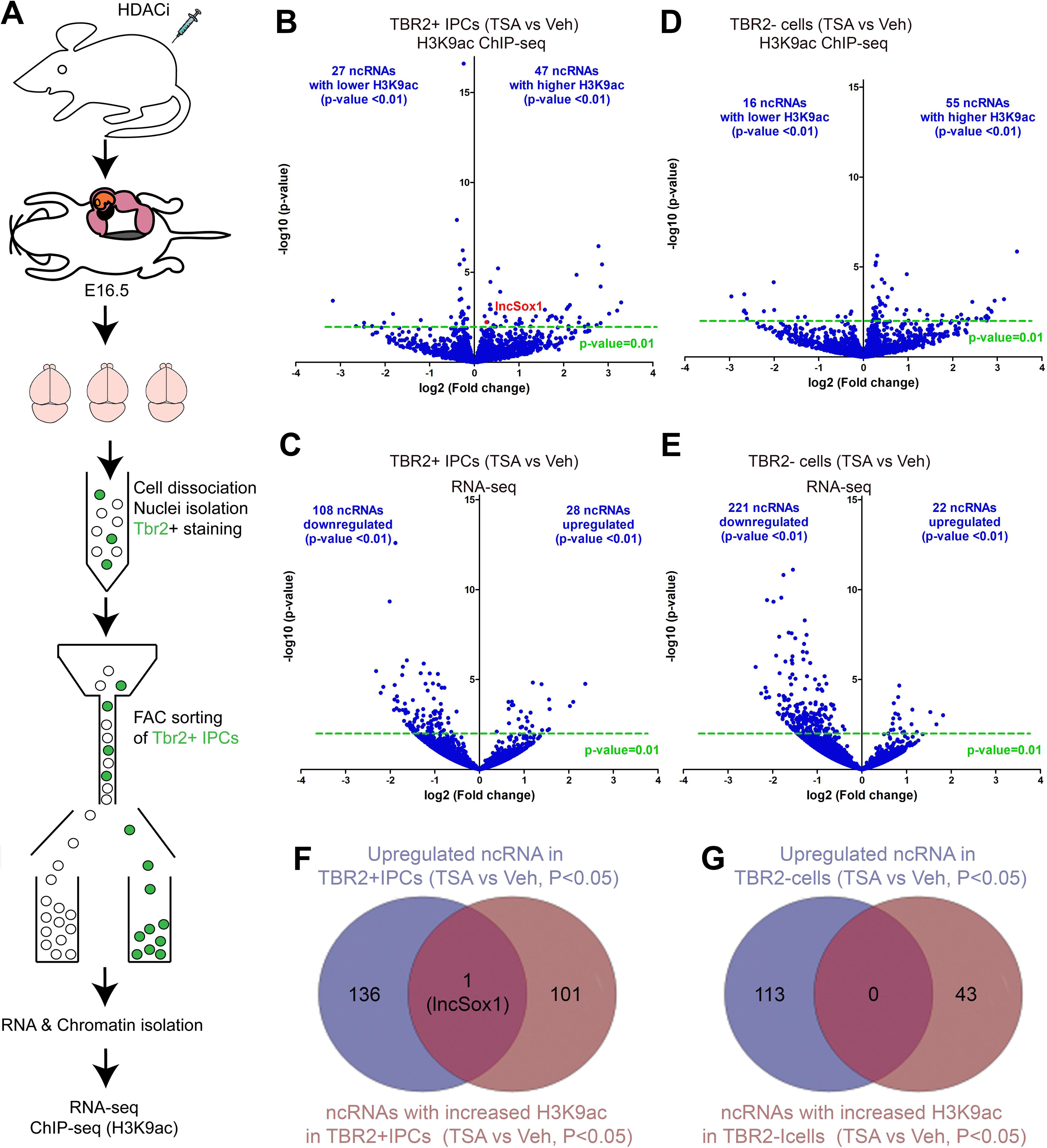
Alteration in the ncRNA milieu in IPCs is a key response to global histone deacetylase inhibition. (A) Schematic overview of the protocol used for profiling the transcriptome and epigenome of IPCs after TSA treatment. (B–E) Volcano plots showing upregulated and downregulated ncRNA genes in terms of their association with the H3K9ac mark (B/D) and expression (C/E) after TSA treatment of IPCs (B/C) and non-IPCs (D/E) compared with their Vehicle-treated counterparts. (F, G) Overlap between upregulated ncRNA genes and ncRNA genes with increased H3K9ac due to TSA-treatment of TBR2+ IPCs (F), and TBR2-IPCs (G). Abbreviation: IPCs, intermediate progenitor cells; TSA, Trichostatin A; HDACi, Histone deacetylation inhibitor.

The TSA-treated E16.5 embryonic brains were harvested and processed for IPC isolation via a FACS protocol based on intranuclear immunostaining of IPCs (Sakib et al., 2021). We used TBR2 antibody to label the IPCs in single-cell suspensions and subsequently isolated the TBR2 positive (TBR2+) cells from non-IPCs [i.e., TBR2 negative (TBR2-) cells] (Fig. 1A) (Sakib et al., 2021; Ulmke et al., 2021). For comparative analysis, the sorted cells were matched such that TBR2+ and TBR2-cells treated with TSA were compared with TBR2+ and TBR2-cells treated with vehicle (Veh), respectively.

To determine how H3K9ac drives IPC generation in the developing cortex, we probed for epigenomic and transcriptomic signatures in IPCs as against non-IPCs with or without TSA treatment. Thus, we performed chromatin immunoprecipitation sequencing (ChIP-seq) and RNA sequencing (RNA-seq) to profile the genome-wide distribution of H3K9ac and H3K9ac-dependent gene expression in the sorted cells. We observed that TSA treatment of the developing brain resulted in notable changes in the expression of both protein-encoding genes (Kerimoglu et al., 2021) and also genes encoding for ncRNAs in both treated IPCs and non-IPCs (Fig. 1B–E). Notably, the TSA treatment of TBR2+ IPCs evoked upregulation of H3K9ac partly related to the loci of 47 ncRNAs as opposed to 27 ncRNAs which displayed lower H3K9ac distribution (*p*-value < 0.01) (Fig. 1B, Table S1). At the transcriptome level, we found that 28 ncRNA transcripts are upregulated, whereas 108 ncRNA transcripts are downregulated upon inhibition of H3 deacetylation in IPCs (*p*-value < 0.01) (Fig. 1C, Fig. S1, and Table S2). On the other hand, 55 and 16 ncRNAs in TBR2-cells displayed high and low H3K9ac, respectively (*p* < 0.01) (Fig. 1D, Table S3). However, our gene expression analysis revealed that 22 ncRNAs are upregulated and 221 ncRNAs are downregulated in such non-IPCs without TSA treatment (*p*-value < 0.01) (Fig. 1E, Table S4). Indeed, some ncRNAs are prominently expressed in non-Tbr2 expressing cell in the presumptive cortical plate (Fig. S1).

To identify potentially direct ncRNA targets of H3K9ac, we compared the upregulated ncRNAs (*p*-value < 0.05) and ncRNAs with increased H3K9ac level (*p*-value < 0.05) in response to TSA treatment in TBR2+ IPCs (Fig. 1F) and in TBR2-cells (Fig. 1G). Interestingly, our comparison revealed lncSox1 as potential downstream factor of H3 acetylation ascribable to the IPC amplification in brain evolution (Fig. 1F). Together, our observations here show that H3K9ac enrichment in IPCs leads to alterations in the ncRNA landscape therein; with an impressive observation that the ncRNA IncSox1 may be directly targeted for upregulation in IPCs following global inhibition of H3 deacetylation in the developing mouse brain.

### 2. TSA treatment increased promoter H3K9ac level and expression of lncSox1 specifically in IPCs

Following a fine screening of the ncRNA output of our ChIP-seq and RNA-seq analyses, we noticeably found lncSox1 to be the only ncRNA with a significantly (*p* < 0.05) high H3K9ac and gene expression upregulation in IPCs due to TSA treatment (Fig. 1F). However, in the case of non-IPCs treated with TSA, there was no specific ncRNA showing a significant (*p* < 0.05) increase in H3K9ac with concurrent gene expression upregulation (Fig. 1F).

As presented in Fig. 1F, lncSox1 expression seems to be particularly upregulated in the sorted IPCs treated with TSA. We further analyzed if there is any element of selectivity in targeting IncSox1 among the pool of ncRNAs with increased expression in TBR2+ IPCs in response to H3K9ac enhancement due to TSA treatment. Indeed, we found lncSox1 as a top upregulated gene in the transcriptome of the TBR2+ IPCs treated with TSA (Fig. 2A). However, whereas the expression of ncRNAs is upregulated in TBR2+ cells as a result of TSA treatment, the expression of lncSox1 is not significantly (*p* < 0.01) increased in non-IPCs in the TSA-treated brain (Fig. 1G, Fig. 2B). Further expression analysis showed that under the condition of TSA treatment there is more than 2-fold increase in IncSox1 expression in TBR2+ IPCs compared with TBR2-cells (Fig. 2C). The observation is consistent with our previous report in which we showed that a small set of genes are specifically upregulated in IPCs due to TSA treatment (Kerimoglu et al., 2021).

**Figure 2.**
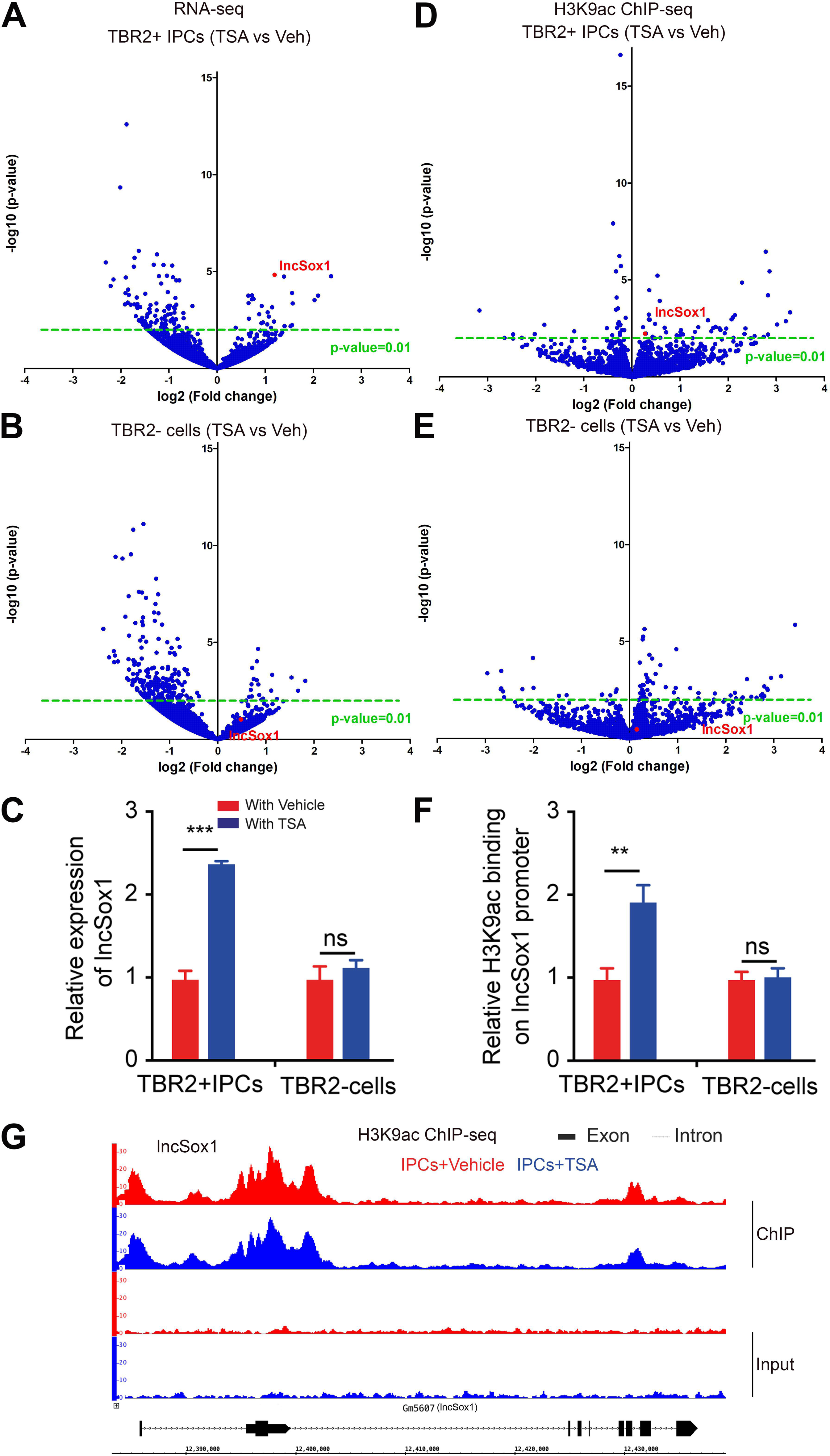
*lncSox1* promoter is a target for TSA-mediated H3K9ac upregulation leading to increased *lncSox1* expression in IPCs. (A–F) Volcano plots showing upregulated and downregulated genes (*p* < 0.01) in the TSA-treated IPC (A/D) and non-IPC (B/E) transcriptome compared with their Vehicle-treated counterparts. lncSox1 has been highlighted in the expression profile as significantly upregulated in (A/D) but with low expression in (B/E). (C, F) Graphs showing relatively increased expression of lncSox1 (C) and H3K9ac binding (F) in TBR2+ cells following TSA treatment compared with control, whereas lncSox1 expression (C) and H3K9ac binding (F) remained unchanged in TBR2-cells upon TSA treatment. (G) Genome browser view of the level and distribution of H3K9ac along the gene body of IncSox1 in TSA-treated (blue) and Vehicle-treated IPCs (red). Input (lower two rows) and distributions after immunoprecipitation (upper two rows) are indicated. Values are presented as means ± SEMs (\*\**p* < 0.001, \*\*\**p* < 0.0001, ns: not significant).

A similar trend in expressing discrepancy was observed for H3K9ac level on IncSox1 promoter following H3K9ac ChIP-seq in TBR2+ IPCs compared with TBR2-cells (Fig. 2D vs Fig. 2E). Accordingly, we found a difference in promoter binding of H3K9ac in TBR2+ IPCs as compared with TBR2-cells, showing almost a 2-fold upregulation in the latter (Fig. 2F). The gene body of *IncSox1* displayed a high level of H3K9ac, particular in the promoter region (Fig. 2G).

Together, our observation here gives an impression of an upregulated gene expression inductive effect of increased H3 acetylation at the promoter of *IncSox1* in TBR2-expressing IPCs following HDAC inhibition by TSA treatment.

### 3. Mouse IPCs have low expression and promoter H3K9ac level of lncSox1 compared with human IPCs

We found in RNA-seq data that the expression of lncSox1 in TBR2-expressing cells in the developing mouse cortex (Fig. 3A/B) is lower than that of human cells (Fig. 3C/D). Among 1428 genes normally downregulated in TBR2+ cells, *lncSox1* ranks as one of the top genes with decreased expression (p < 0.01, |FC > 1.0|) in the mouse cortex (Fig. 3A). However, TBR2+ cells in human cortex displayed a high level of *lncSox1* expression, hence it was found among the 2053 genes upregulated (p < 0.01, |FC > 1.0|) in the human TBR2-expressing cells (Fig. 3C). Interestingly, the relative amount of *lncSox1* transcripts in TBR2-cells is much higher than that found in TBR2+ IPCs in mouse (Fig. 3B), whereas the converse is true for that of human (Fig. 3D). Indeed, expression analysis in the developing (E14.5) mouse cortex shows *lncSox1* expression is concentrated in the ventricular zone (VZ) of the neocortex, where majority of cells are radial glial cells (RGCs) (Fig. S2A). Along this line, scRNA-seq analysis (Telley et al., 2016) revealed that *lncSox1* is expressed in non-IPCs in mouse cortex with its highest level was seen in RGCs and early-born neurons (EN) (Fig. S2B). On the other hand, IPCs and late-born neurons (LN) are very low in *lncSox1* expression (Fig. S2B).

**Figure 3.**
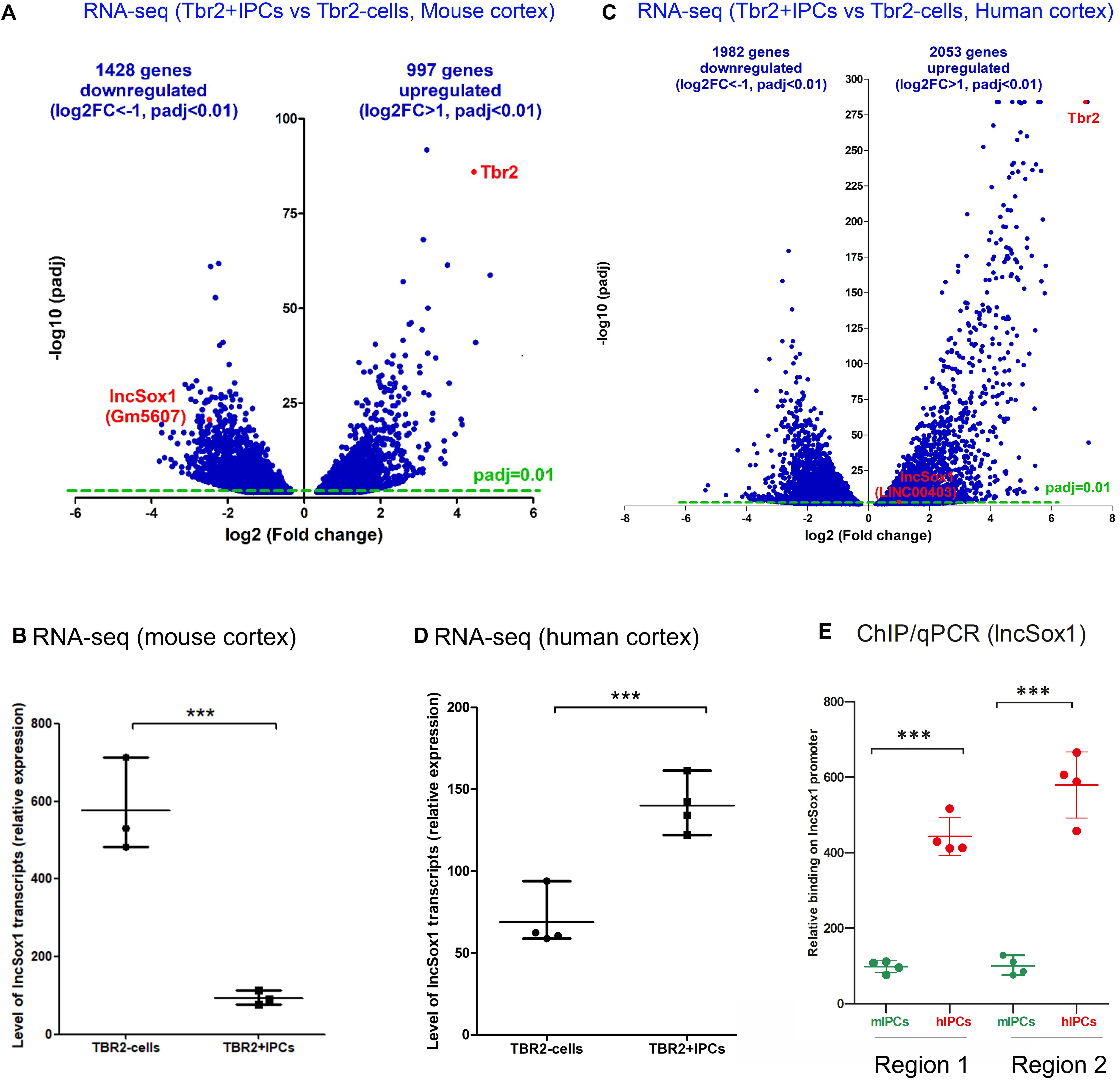
Low baseline level of H3K9ac at *lncSox1* promoter and *lncSox1* expression in IPCs. (A) Volcano plot showing upregulated and downregulated genes (*p* < 0.01; FC > 1.0) in the developing mouse cortex transcriptome. The gene expression profile shows two clusters of genes: IPC and non-IPC genes, with *lncSox1* indicated as one of the upregulated non-IPCs genes. (B) Graph showing relative expression of *lncSox1* in TBR2-cells compared with TBR2+ cells. (C) *In situ* hybridization micrograph showing the expression of *lncSox1* in the E14.5 mouse brain. (D) Image showing the expression profile of lncSox1 in various cell-types in the cortex based on single-cell (sc)RNA-seq. (E) Volcano plot showing genes with high and low levels of H3K9ac following H3K9ac ChIP-seq in TBR2+ cells compared with TBR2-cells. *lncSox1* has been highlighted as one of the genes with low level of H3K9ac in TBR2+ cells. (F) Graph showing relative expression of H3K9ac in TBR2+ cells compared with TBR2-cells. Values are presented as means ± SEMs (***P < 0.0001).

Of note, the low expression of lncSox1 correlates with a low promoter H3K9ac signal at the *lncSox1* locus in the TBR2+ IPCs found in the mouse cortex (Fig. S2C). ChIP and qPCR experiments comparing the level of H3K9ac in TBR2+ IPCs indicated a significantly lower level of *lncSox1* promoter H3K9ac in mouse IPCs than that of human IPCs (Fig. 3E, Fig. S2C).

The finding indicates that the normal differential expression of lncSox1 in TBR2+ IPCs in mouse and human is linked to a difference in H3K9ac enrichment at the *lncSox1* promoter region, which is high in human IPCs and likely responsible for the lncSox1 upregulation. Thus, this observation is a plausible explanation for why there is increased lncSox1 expression in TBR2+ IPCs following induction of high H3K9ac as a result of TSA treatment (Fig. 1F, Fig. 2A-F).

### 4. Targeted H3K9 acetylation increases the promoter H3K9ac and expression of lncSox1

To validate that increased H3K9 acetylation specifically at the *lncSox1* locus is necessary for promoting the expression of lncSox1, we employed a CRISPR/dCas9-based system (Albert et al., 2017; Kerimoglu et al., 2021) which allowed targeted depositions of H3K9ac mark at the *lncSox1* locus (Fig. 4A). We generated plasmid constructs (gLncSox1-dCas9-KAT2A-T2A-eGFP) bearing DNA sequences that encoded a guide RNA (gRNA), and dCas9 (a modified Cas9 without nuclease function) fused to the acetyltransferase KAT2A, and GFP as a fluorescent reporter (Fig. 4B). Thus, the approach constitutes a targeted epigenome editing leading to the remodeling of the H3K9ac landscape at the promoter region of *lncSox1*.

**Figure 4.**
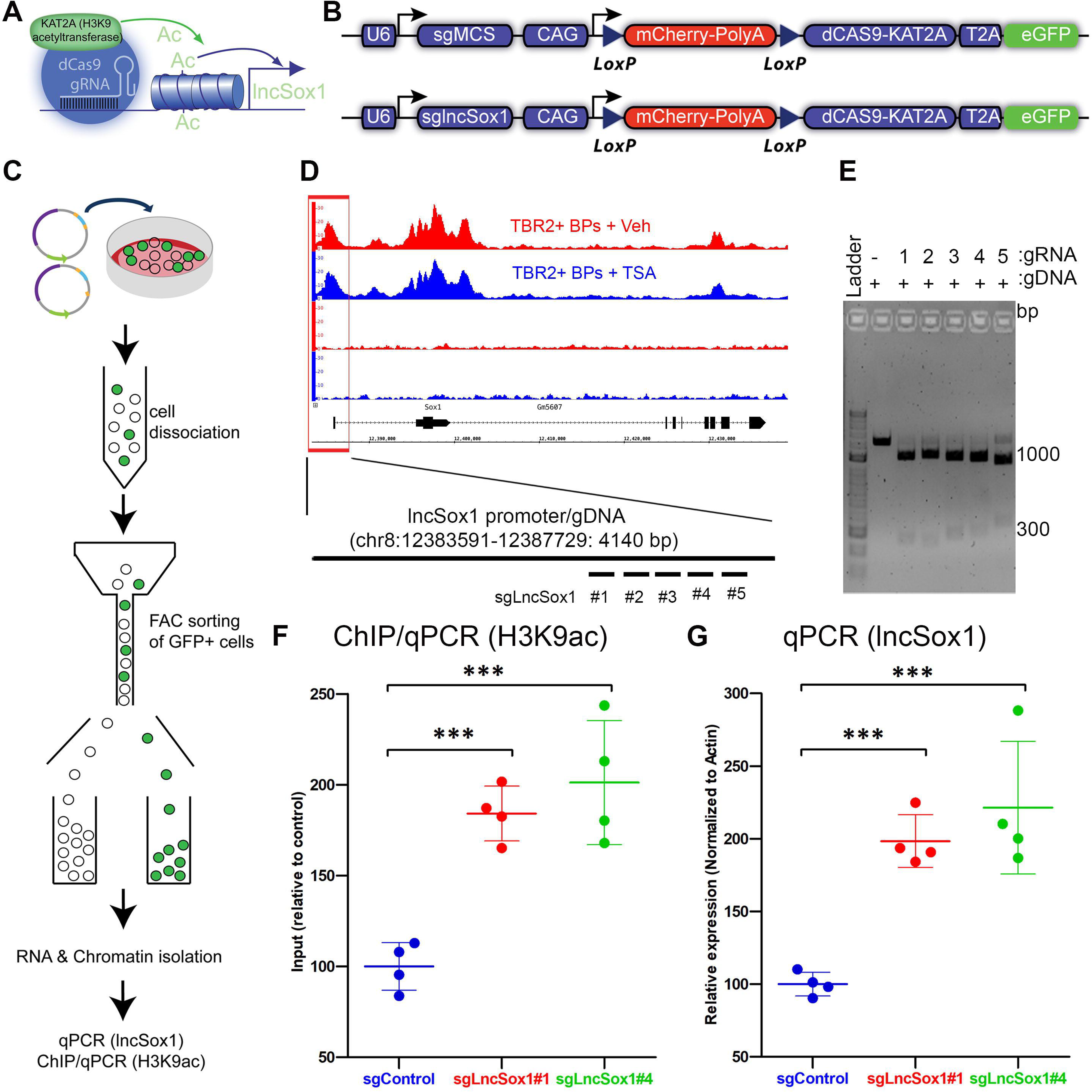
Editing H3K9 acetylation specifically at *lncSox1* promoter has implication for enhanced lncSox1 expression. (A) Illustrative overview of the CRISPR/dCas9-mediated installation of H3K9ac at the lncSox1 promoter to augment lncSox1 expression. (B) Schema showing CRISPR/dCas9 constructs used for validating the phenotypic effect of H3 acetylation at the lncSox1 promoter. (C) Diagrammatic representation of the neuro2A cell culture system used for testing the CRISPR/dCas9 constructs for editing H3K9ac at lncSox1 promoter followed by FAC sorting of transfected (GFP+), and isolation of chromatin and RNA for the quantification of H3K9ac and IncSox1 expression, respectively. (D) Genome browser view showing the level and distribution of H3K9ac along the gene body of IncSox1 in TSA-treated IPC (blue) and Vehicle-treated IPCs (red). Input (lower two rows) and distributions after immunoprecipitation (upper two rows) are indicated. The various tested gRNAs targeting different regions of IncSox1 promoter for H3K9ac editing are indicated. (E) Image of agarose gel showing the cutting efficiency of each gRNA-Cas9 complex tested on a 4140-bp-long PCR product of IncSox1 promoter region. (F, H) Graphs showing increased level of H3K9ac at IncSox1 promoter (F), and elevation in the expression of IncSox1 (G) in the cultured Neuro2A transfected with the gLncSox1#1 and #4. Values are presented as means ± SEMs (***p < 0.0001).

To target H3K9 acetylation at *lncSox1* promoter in a cell type-specific manner, we employed the Cre/loxP system in our epigenome editing construct, in which an mCherry-polyA sequence flanked by loxP sites was placed in front of the dCas9-KAT2A sequence (Fig. 4B). Using sorted cultured neurons (Neuro2A), we validated several designed gRNAs capable of targeting the *lncSox1* promoter. Multiple lncSox1 sgRNAs (sglncSox1 #1–5), were used to target the *lncSox1* promoter for acetylation by the H3K9ac writer KAT2A (Fig. 4C-E).

Upon sorting the Neuro2A cells co-transfected with the H3K9ac-installing construct together with pCAG-Cre plasmid (i.e., GFP-expressing cells), and performing ChIP-qPCR experiment (Fig. 4C), we observed a significant (*p* < 0.0001) elevation in the level of H3K9ac at the *lncSox1* promoter due to deposition of H3K9ac under the gLncSox1#1 and gLncSox1#2 as compared with the control construct (gControl) (Fig. 4F). Additionally, based on qPCR analysis, we found an increased level of lncSox1 in the isolated cells transfected with the generated H3K9ac editing constructs (Fig. 4G). These results indicate validation of the CRISPR/Cas9-mediated epigenome editing system as capable of H3K9ac deposition at the promoter of *lncSox1*, which resulted in increased expression of lncSox1 in neural cells.

### 5. Targeted H3K9 acetylation at *lncSox1* promoter in IPCs allows investigation of the associated phenotype

To achieve epigenome editing-based regulation of H3K9ac levels in IPCs *in vivo*, we delivered the constructs into tamoxifen (TAM)-treated TBR2/Eomes-CreER mouse embryos (Pimeisl et al., 2013) by IUE at E13.5 (Fig. 5A/B). Cre recombinase is driven by the IPC-specific TBR2 regulatory elements, which switch expression from mCherry to dCAS9-KAT2A and eGFP specifically in IPCs and their progenies (Fig. 5A/B). To validate our cell type-specific epigenome editing system, we examined the location of mCherry+, and eGFP+ cells and their co-expression with the IPC marker TBR2.

**Figure 5:**
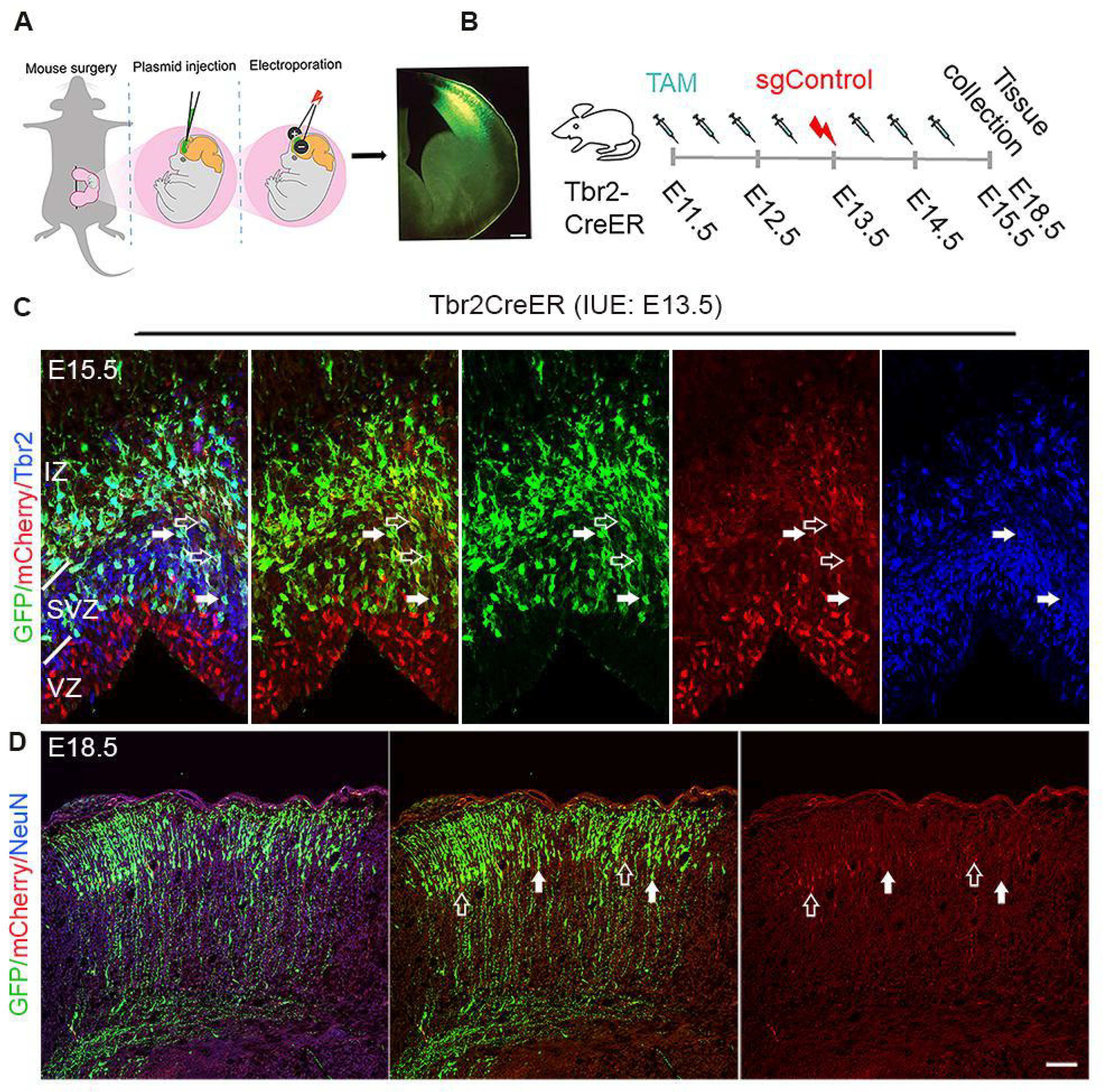
Cortical phenotype analysis following editing of H3K9ac at the promoter region of *IncSox1*. (A) Schematic overview of *in utero* electroporation of the embryonic mouse cortex with sgLncSox1 and Tbr2CreER plasmids. (B) Illustration of treatment of mouse embryos with tamoxifen (TAM) and Tbr2CreER via scheduled intraperitoneal injections. (C, D) Immunohistochemistry micrographs of the E15.5 (C) and E18.5 cortex stained for the indicated markers following the *in utero* electroporation-mediated H3K9ac editing at the promoter region of *IncSox1*. White arrows in (C, D) point to mCherry-non-labelled cortical cells exposed to CreER and with faint or no TBR2 labelling (C) in the E15.5 cortical germinal zone (C) and E18.5 cortical plate (D). Empty arrows in (C, D) point to mCherry-labelled cells with no or faint TBR2 expressing (in C) in the E15.5 cortical germinal zone (C) and E18.5 cortical plate (D). Abbreviations: VZ, ventricular zone; SVZ, subventricular zone; IZ, intermediate zone. Scale bars = 50 µm.

At E15.5, the majority of cells expressing high level of mCherry are found in TBR2-cells in the VZ (Fig. 5C/D). As expected, most of the eGFP+ cells were found in TBR2+ cells in SVZ, and possibly neuronal progenies of IPCs in IZ/CP (Fig.5C, filled arrows). At E15.5, we noted that many eGFP+ cells in SVZ/IZ also expressed mCherry at a low level, possibly due to mCherry proteins stability over few days (Fig. 5C, empty arrow). Indeed, all eGFP and mCherry do not co-express in the cortex at a later stage at E18.5 (Fig. 5D, arrows). Thus, the established cell type-specific epigenome editing system allows us to add the epigenetic mark H3K9ac onto *lncSox1* promoter specifically in IPCs and their progenies in developing mouse cortex.

### 6. IPC-specific H3K9 acetyl editing at *lncSox1* promoter promotes IPC proliferation and indirect neurogenesis

To determine the impact of increasing the expression of lncSox1 in IPCs through targeted increase in the promoter level of H3K9ac, we immunohistochemically analyzed the developing mouse cortex treated with our epigenome editing tool. At E14.5, the cortex electroporated with gLncSox1 plasmid displayed a greater ratio of TBR2+/GFP+ cells in the SVZ per mCherry+/TBR2-cells in the VZ than that in the control plasmid-injected cortex (Fig. S3A), suggesting TBR2+ IPC population was expanded upon gLncSox1 expression. We also analyzed the proliferative capacity of TBR2+ IPCs in the developing cortex (E15.5) following the lncSox1 expression induction paradigm (Fig. 6A-D). An assessment of the proportion of TBR2+/BrdU+ (IPCs in S-phase) and TBR2+/pHH3+ (IPCs in M-phase) cells among the GFP-labelled TBR2+ IPCs in gControl- and gLncSox1 construct-injected cortices enabled us to determine if the H3K9ac level alterations at the *lncSox1* promoter contributed to the proliferation of IPCs. Remarkably, the increase in promoter H3K9ac level of lncSox1 by our epigenome editing tool augmented the amount (percentage) of basal mitosis (BrdU+ or TBR2+/pHH3/GFP+) among targeted IPCs (TBR2+/GFP+; Fig. 6A-D).

**Figure 6.**
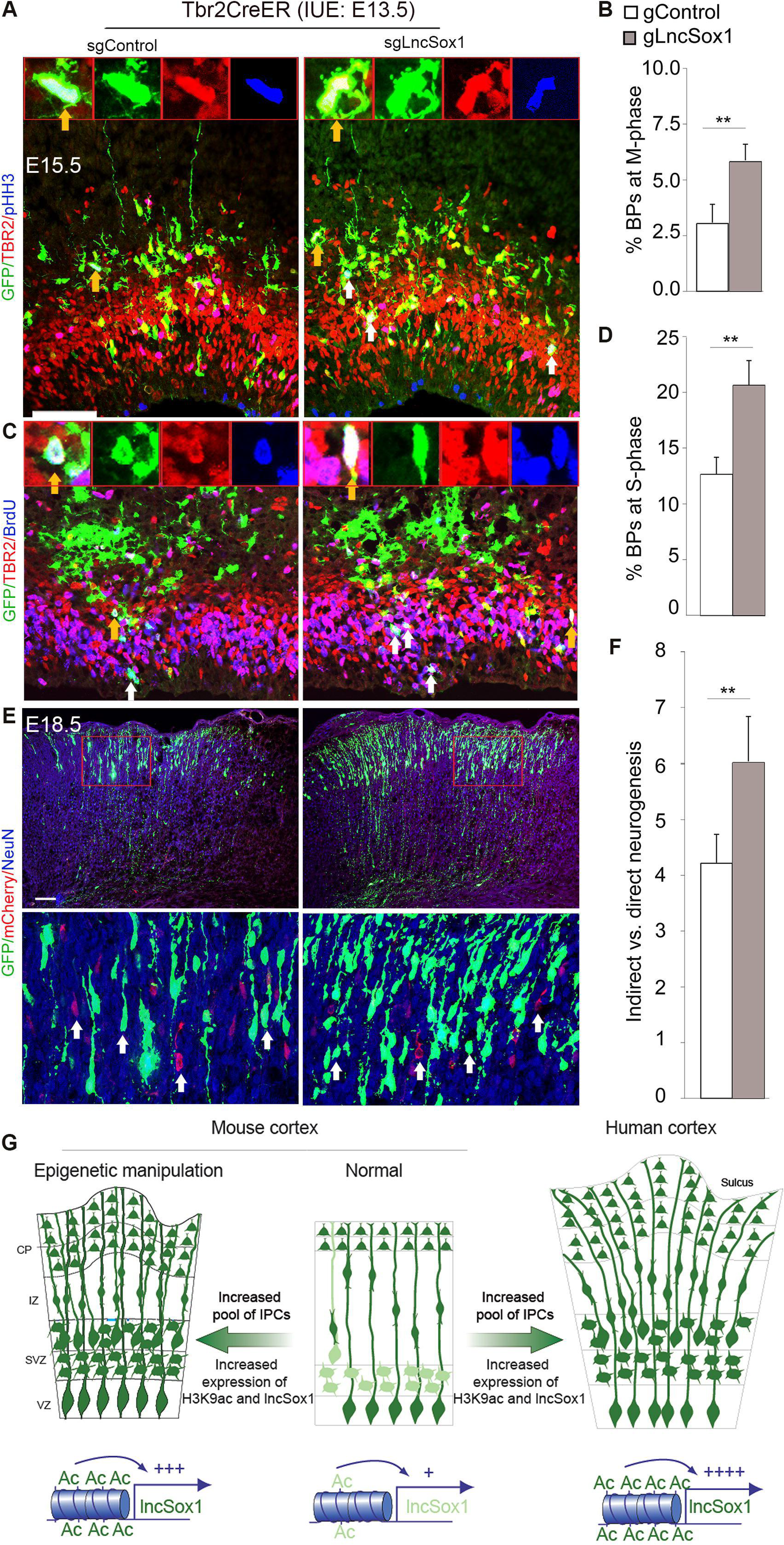
Targeted H3K9 acetylation at *lncSox1* promoter encourages IPC proliferation and neurogenesis. (A) Micrographs showing immunostaining analysis of pHH3, TBR2, GFP in the E15.5 mouse cortex electroporated at E13.5 with sgLncSox1 and sgControl plasmids. Inserts are high magnifications of cells stained with the indicated markers and display M-phase status. Yellow arrows point to TBR2+ cells with mitotic figures and white arrows point to those without mitotic figures. (B) Statistical quantification of proliferating TBR2-expressing cells in M-phase in the sgLncSox1- and sgControl-treated E15.5 cortex. (C) Micrographs showing immunostaining analysis of BrdU, TBR2, GFP in the E15.5 mouse cortex electroporated with sgLncSox1 and sgControl plasmids. Inserts are high magnifications of cells stained with the indicated markers and display S-phase status. Yellow arrows point to TBR2+ cells with active DNA replication and white arrows point to those without active DNA replication. (D) Statistical quantification of proliferating TBR2-expressing cells in S-phase in the sgLncSox1- and sgControl-treated E15.5 cortex. (E) Micrographs at low (upper panels) and high (lower panels) magnifications showing GFP, mCherry, and NeuN immunostaining in the E18.5 mouse cortex electroporated at E13.5 with sgLncSox1 and sgControl plasmids. Red rectangle demarcates zoomed area of the cortical plate shown in the lower panel micrographs for quantitative analysis. White arrows point to mCherry/NeuN cells. Their lack of GFP expression indicate they are derived from non-TBR2-expressing cells or direct neurogenesis. Cells with GFP/mCherry/NeuN staining are derived from TBR2+ cells via indirect neurogenesis. (F) Statistical quantification of the proportion of cells undergoing direct or indirect neurogenesis in the sgLncSox1- and sgControl-treated E18.5 cortex. Values are presented as mean ± SEM (\*\**p* < 0.01). Scale bars = 50 μm. (G) Schema illustrating how lncSox1 expression in the normal mouse cortex is augmented by its promoter region H3K9ac enrichment leading to intermediate progenitor pool expansion, which has implication for expansion and folding of the epigenetically manipulated mouse cortex. The outcome of such epigenetic manipulation phenocopies the human cortical with possibly evolutionary basis. Abbreviations: Ac, H3K9ac mark; IPCs, intermediate progenitor; VZ, ventricular zone; SVZ, subventricular zone; IZ, intermediate zone; CP, cortical plate.

During corticogenesis, RGCs are actively involved in direct neurogenesis mode and undergo indirect neurogenesis via IPCs. To exclude the possibility that our IPC-specific H3K9ac editing system does not influence direct neurogenesis, we also used the same approach to deliver the expression plasmids in the E13.5 cortex. Given that RGCs undergo only one division in 24 hr at mid-gestation (Noctor et al., 2004), we collected tissue at E14.5 and performed double immunostaining for mCherry and neuronal markers Tuj or HuCD to quantify the neuronal production directly from mCherry+ RGCs. In the control, electroporated (mCherry+) cells were located in the SVZ/IZ and some coexpressed Tuj1/HuCD (22% ± 2.6% of all mCherry+ cells), representing differentiated neurons generated directly from RGCs (Fig. S3B-E). Compared to control, the gLncSox1 plasmid-electroporated cortex has a similar proportion of mCherry+/Tuj1/HuCD+ neurons among the targeted mCherry+ cells, indicating that the IPC-specific gLncSox1 activation does not affect the direct neurogenesis.

To examine whether the IPC-specific gLncSox1 overexpression promotes indirect neurogenesis, we quantified the number of GFP+/NeuN+ or Satb2+ neurons in the gLncSox1 or gControl plasmids-electroporated cortices at E18.5. Because the direct neurogenesis is not different between the gLncSox1 and empty vector plasmids (EV)-electroporated cortices, we used mCherry+/NeuN+ or mCherry+/Satb2+ neurons as internal control for our comparison (Fig. 6E). Our comparative analysis indicated that the gLncSox1-electroporated cortices have a greater ratio of GFP+ neurons per mCherry+neurons than that of EV plasmids-electroporated cortices and indicative of increased indirect neurogenesis (Fig. 6F).

Because we found that apoptosis in the VZ/SVZ, cortical layers, and direct neurogenesis were not significantly different between gLncSox1 gain-of-function effect and that of control, these findings indicate that IPC-specific activation of gLncSox1 promotes cortical neurogenesis via increasing IPC proliferation.

## DISCUSSION

Brain evolution can be partly described on the basis of the adoption and/or operationalization of new and advanced regulatory mechanisms or the hierarchical prominence of such mechanisms in the course of evolution. Notably, proliferative neurogenic cells such as the TBR2-expressing IPCs are critical determinants of brain development, and there is interspecies variation in the abundance of TBR2+ IPCs in the developing brain. Therefore, an understanding of the molecular factors driving this variation would afford an *evo-devo* insight into brain organogenesis and expansion. For instance, emergence of the evidence that H3 acetylation in basal progenitor cells is a key mechanism underlying cortical developing (Kerimoglu et al., 2021) have provoked questions about how the neuroepigenome contributes to the evolutionary expansion of the mammalian brain. However, it is challenging to target specific epigenetic events to study their role in specific population of cells. We attempted answering such questions by seeking to unravel the precise regulatory epigenetic mechanisms which may drive the proliferative capacity of IPCs in the developing cortex, with plausibly implication for brain evolution. Our quest brought into focus the exploration of targeted gene regulation in a purified population of neural cells. This led to the concept of combining the epigenome editing strategy and FACS-mediated cell isolation protocol to cell type-specifically probe for the epigenetic factor(s) involved in IPC biogenesis during cortical expansion. The advantage offered by epigenome editing is the effective manipulation of the genomic phenotype without direct interference of the nucleotide sequence in the gene(s) of interest.

We went about our neuroepigenome editing objective by initially treating the developing mouse cortex with TSA to chemically inhibit histone deacetylation, which is essentially an indirect approach to upregulating histone acetylation (Kerimoglu et al., 2021). We then isolated IPCs from the TSA-treated brains. The use of the intranuclear immunostaining FACS method for purifying IPCs was to ensure comprehensive isolation of TBR2-expressing IPCs in the developing cortex (Sakib et al., 2021). Next, we examined the IPC transcriptome for key epigenetic changes in response to the histone acetylation changes. The RNA-seq analysis and Chip-seq experimentation were used to probe for epigenetic changes which distinguish TBR2+ cells from TBR2-cells in the context of response to increase in histone acetylation. Our observation of overt alteration in ncRNA expression in the sorted TSA-treated TBR2+ IPCs led to the identification of lncSox1 expression as a major downstream effect of histone acetylation upregulation in IPCs. This outcome is consistent with the observation that targeted increase in histone acetylation at promoters and enhancers is sufficient to enhance gene expression (Hilton et al., 2015). Indeed, by using our epigenome editing tool, we were able to efficiently deposit H3K9ac at the promoter of *lncSox1* which resulted in its increased expression. The strategy of targeting different promoter regions of *lncSox1* gene with multiple *lncSox1* gRNAs was to ensure efficient targeting leading to adequate achievement of the desired manipulative effect, i.e., H3K9ac addition at the promoter region. This strategy has gained popularity because it helps in achieving robust activation of genes (Cheng et al., 2013; Gilbert et al., 2013; Hilton et al., 2015; Konermann et al., 2015; Liu et al., 2016; Mali et al., 2013; Perez-Pinera et al., 2013; Qi et al., 2013).

Application of the CRISPR-dCas9 platform to effect durable gene expression regulation in the neuroepigenome is fast becoming one of the most reliable and highly efficient strategies for identifying the specific contribution of particular epigenetic factors, including chromatin marks, in neurodevelopment (Yim et al., 2020). A fascinating feature of the CRISPR-dCas9-mediated epigenome editing is the programmability of the gRNA component, which allows targeted manipulation at specific gene loci (Zalatan et al., 2015). In other to validate the specificity of our epigenome editing system, we performed RNA-seq and ChIP-seq as commonly done (reviewed in (Yim et al., 2020)). Our gLncSox1 was able to efficiently target the acetyltransferase KAT2A to the promoter region of *lncSox1* gene for H3K9ac installation leading to the inductive upregulation of lncSox1 expression. The use of TBR2-Cre for the activation of the CRISPR-dCas9 construct ensured the cell type-specific targeting of TBR2-expressing IPCs *in vivo*. Other previously reported ways of increasing the efficient of an epigenome editing machinery worth considering for improving CRISPR-dCas9-dependent epigenome engineering specificity include (i) using shorter (18–19 base pairs) gRNAs to increase binding complementarity (Fu et al., 2014), (ii) point mutating a positively charged domain in the CRISPR-Cas9 structure to reduce gRNA off-targeting (Slaymaker et al., 2016), and (iii) modifying the secondary structure of gRNAs by adding a hairpin loop to inhibit Cas9 off-target binding (Kocak et al., 2019).

To validate our epigenome editing scheme, we sought to analyze the outcome of editing H3K9ac at the promoter region of *lncSox1* in IPCs during cortical development. The observed striking consequence of increase in the IPC pool is reasonably linked to the upregulation of *lncSox1* following the H3K9ac enhancement manipulation. Our finding that the lncSox1-enriched IPCs exhibit enhanced proliferative capacity explains the demonstrable increase in the population of the IPCs and the resultant increase in neurogenesis. This observed effect of our targeted epigenome editing experiment in the developing mouse cortex offers opportunity to uncover novel mechanisms in specific population of cells which can elucidate developmental processes during cortical growth.

Put together, by means of our epigenome editing paradigm, the histone mark H3K9ac has been identified as an epigenetic signature of IPCs mediated by selected ncRNAs such as *lncSox1*, which is likely evoked as an evolutionary requirement for increasing the neural progenitor pool in the mammalian developing cortex (Fig. 5G). The CRISPR-dCas9-based epigenome editing method has thus been validated as an efficient tool for robust interrogation of endogenous gene function in the neuroepigenome. The method is therefore suitable for studying specific epigenetic events at defined gene loci in specific cell types to afford precise description of their contribution to cellular processes *in vivo*.

## METHOD DETAILS

### Transgenic mice and *in utero* electroporation (IUE)

Tbr2/EomesCreER mice (Pimeisl et al., 2013) were maintained in a C57BL6/J background. Animals were handled according to the German Animal Protection Law. *In utero* electroporation was done as previously described (Tuoc and Stoykova 2008; Tuoc et al. 2013).

### Plasmids

Plasmids used in this study: pCIG2-Cre-ires-eGFP (a gift from Prof Francois Guillemot, NIMR London), sgLncSox1#1#2-LoxP-mCherry-PolyA-LoxP-dCas9-KAT2A-T2A-eGFP (this study).

### Antibodies

Monoclonal (mAb) and polyclonal (pAb) primary antibodies were commercially sourced and listed as follows: GFP chick pAb (1:400; Abcam), mCherry/RFP pAb (1:1000, biomol/Rockland), mCherry/RFP mAb (1:1000, biomol/Rockland), NeuN mouse mAb (1:200, Chemicon), HuCD mouse mAb (1:20; Invitrogen), Pax6 rabbit pAb (1:200; Covance), Pax6 mouse mAb (1:100; Developmental Studies Hybridoma Bank), TBR2 rat 923 mAb (1:200; eBioscience), TBR2 rabbit pAb (1:200; Abcam), H3K9ac pAb (ChIP-seq, Millipore), pHH3 mAb (1:50; Cell Signaling), BrdU rat pAb (1:100; Abcam), BrdU mouse mAb (1:40; CalTag), Tuj mAb (1:200; Chemicon). Secondary antibodies from various species used are Alexa (488, 568, 594 and 647)-conjugated IgG (1:400; Molecular Probes).

### Mouse treatment with HDAC inhibitors (HDACi) (Kerimoglu et al., 2021)

Trichostatin A (TSA) solution was at a concentration of 100µg/ml by dissolving in vehicle (8% ethanol in 1xPBS). Pregnant mice from E12.5 dpc were treated twice daily with either 150µl of 100 µg/ml TSA solution or vehicle via intraperitoneal injection and sacrificed at various stages of embryonic development indicated in the text.

### TBR2-stained nuclei and cell sorting from embryonic mouse cortex (Kerimoglu et al., 2021)

#### Sorting TBR2+ nuclei for ChIP-Seq

The protocol was adopted from Halder and colleagues (Halder et al., 2016). Unless stated otherwise, steps were done on ice or at 4°C. Replicates of CD1 embryonic cortices (pooled from 5 pubs) were briefly homogenized with plastic pestles in 1.5ml tubes in low-sucrose buffer (320 mM Sucrose, 5mM CaCl2, 5 mM MgAc2, 0.1 mM Ethylenediamine tetraacetic acid (EDTA), 10mM HEPES pH 8, 0.1% Triton X-100, 1mM DTT, supplemented with Roche protease inhibitor cocktail). Incubation with 1% formaldehyde for 10 mins at room temperature was used for crosslinking. Excess formaldehyde in solution was neutralized with 125mM Glycine under same incubation condition for 5 mins. The mixture was centrifuged at 2000xg for 3 mins, the nuclear pellet was resuspended in low-sucrose buffer and mechanical homogenized further with a homogenizer (IKA Ultraturax). The mixture was sedimented in high-sucrose buffer (1000mM Sucrose, 3mM Magnesium acetate, 10mM HEPES pH 8, 1 mM DTT, protease inhibitor) by spinning in oak-ridge tubes at 3200xg for 10 mins in a swinging bucket rotor centrifuge. This effectively eliminates myelin. The supernatant was carefully decanted, and nuclei were collected in 2ml DNA-low bind microfuge tubes following 3 mins centrifugation at 2000xg. Excess sucrose was removed, and the pellet of nuclei resuspended with 500uL PBTB buffer (PBS-0.2% Tween-1% BSA buffer, with protease inhibitor). Application of anti-TBR2 for 1 hour was used to achieve Tbr2 staining followed by washing of nuclei and resuspension with 500µl PBTB. Tbr2 nuclei sorting was used to performing using FACSAria III. Samples with no antibody were used as negative control for gating. Sorted nuclei were collected into 15mL falcon tubes coated with PBTB and briefly centrifuged for the Tbr2 stained and unstained nuclei to pellet. Liquid nitrogen was used to flash freeze the Sorted nuclei pellets which was subsequently stored at −80°C for later use in ChIP experiment.

#### Sorting TBR2+ nuclei for RNA isolation and Sequencing

Freshly dissected CD1 embryonic cortices (pooled from 5 pubs) were collected and immersed in RNAlater solution contained in a microfuge tube, which was kept at 4°C for not less than 24 hours. Two washes with 1x RNAse free PBS were enough to remove excess RNAlater solution from the cortices. Incubation steps were performed on ice and centrifugation at 500xg in 4°C, where not specified. The cortical tissues were homogenized with 500 uL lysis buffer (Sigma, NUC101) using 30-45 plastic pestles agitations. The homogenate was made to 2 mL by adding lysis buffer. Incubation was done for 7 mins after which the lysate was centrifuged for 5 mins and nuclei pellet resuspended in 2ml lysis buffer. The lysate was incubated for 7 mins, filtered into a new 2ml tube using a 40µm filter, and the supernatant removed by centrifugation. The resultant nuclei pellet was washed with 1800ul nuclei suspension buffer (NSB), centrifuged, and resuspended in 500uL NSB. The NSB was prepared as follows: 0.5% RNAse-free BSA (Millipore), 1:200 RNaseIN plus RNAse inhibitor (Promega), protease inhibitor diluted into RNAse free PBS (Invitrogen). Nuclei were stained for Tbr2 by applying anti-TBR2 Alexa 488 conjugated antibody for an hour. Stained nuclei were NSB-washed and resuspended in NSB. Tbr2 stained/unstained nuclei were then sorted into NSB coated falcon tubes for ChIPseq as described in the section above. For RNA isolation, Trizol LS solution was added to nuclei collected by brief centrifugation. This was followed by chloroform addition for nuclei solubilization, and 15 mins centrifugation at 120000xg. Zymo RNA clean & concentrator-5 kit then was used to purified RNA from the collected aqueous phase. By means of an RNA library kit (Takara SMART-Seq v4 Ultra Low Input RNA kit) mRNA-seq libraries were prepared using 1ng of RNA purified from the isolated Tbr2-labelled nuclei. Sequencing of the libraries was done in in Illumina Hiseq 2000 machine.

### Chromatin immunoprecipitation (ChIP)

The ChIP experiment with H3K9ac antibody (Millipore 07-352) was conducted according to our previously described protocol (Narayanan et al., 2015). Immunoprecipitated DNA was analyzed either by qPCR or sequencing, in which Illumina sequencing libraries were generated with the DNA library preparation kit NEBNext Ultra II. Illumina Hiseq 2000 machine was then used to sequence the libraries.

### ChIP and RNA sequencing data analyses

Next generation sequencing and data analyses were performed according to steps described in our previous work (Kerimoglu et al., 2021). RNA-seq was performed and analyzed using our routinely established protocol described in our previous work (Narayanan et al., 2015; Nguyen et al., 2018).

### Identification of lnRNAs enriched in IPCs

The IPC genes with significant expression changes (log2FoldChange > 1.0, p-value < 0.01) were analyzed with the MGI gene nomenclature tool available at (http://www.informatics.jax.org/batch). List of genes which encode for lnRNAs proteins was extracted from each other.

### Histochemistry validation

Immunohistochemistry (IHC) and *in situ* hybridization (ISH) were performed as previously described (Bachmann et al., 2016; Tuoc et al., 2013; Visel et al., 2004). To eliminate fluorescent signal of mCherry in triple IHC analysis, the tissue was treated with HCl.

### qRT-PCR analyses

Western blot analyses and qRT-PCR were performed based on our previously described protocols (Tuoc and Stoykova, 2008). Primers used are shown in Table S6 and the RT² Profiler PCR Array profiles (Qiagen).

### Microscopy and statistical analysis

Histological images were obtained using confocal fluorescence microscopy (TCS SP5, Leica) and analyzed with Axio Imager M2 (Zeiss) with a Neurolucida system. Further processing of micrographs was done using Adobe Photoshop. Quantities were averaged from a minimum of three biological replicates. Statistical analyses for histological experiments are detailed in in Table S10.

### Epigenome editing

#### Design epigenome editing constructs

The backbone vector (U6-sgMCS-CAG-LoxP-mCherry-PolyA-LoxP-dCas9-KAT2A-T2A-eGFP) was commercially synthesized by GenScript. gRNA sequences targeting specific mouse LncSox1 promoter region (chr8:12383591-12387729) was designed using the software ‘Genious’, and inserted into a T7 promoter-based vector.

#### Testing for LncSox1 sgRNAs, quantification of H3K9ac level at lncSox1 promoter and expression of LncSox1 in sgLncSox1-CAG-LoxP-mCherry-PolyA-LoxP-dCas9-KAT2A-T2A-eGFP-transfected Neuro2A

Synthesized sgRNAs were test *in vitro*. The PCR-produced sgRNA template used for the sgRNA synthesis carried the T7 promoter added by the gRNA sequence and the sharped gRNA scaffold. The *in vitro* transcription was done using the transcription kit MEGAscript T7 and following the manufacture’s protocol. Cas9 protein was sourced from IDT (#1074182). The cutting efficiency of Cas9/gRNAs complexes was tested *in vitro* using a IPC PCR product from lncSox1 promoter. Detailed description of the protocol used for the *in vitro* testing is available in our previous work (Kerimoglu et al., 2021). After *in vitro* test, the sgLncSox1(#1, GTGTCTCGAACTCGCGCGCGG) and sgLncSox1(#4, GTGTGTGCCGAACGAGGAGCA) sequences were selected, synthesized and cloned into the backbone vector (U6-sgMCS-CAG-LoxP-mCherry-PolyA-LoxP-dCas9-KAT2A-T2A-eGFP) to generate corresponding plasmids (sgLncSox1#1, sgLncSox1#4). The vectors were sequenced for confirmation. Neuro2A cells were transfected with 12 µg of parental vector pCAG-Cre-ires-eGFP, or/and sgLncSox #1, sgLncSox #4 using lipofectamine 2000 reagent (Thermofisher) and cultured on 10 cm dishes. The cultured cells were collected 3 days after transfection, and FACS analyzed for qPCR and ChIP/qPCR analyses.

## DATA AND CODE AVAILABILITY

RNA-seq and ChIP-seq data have been deposited at the GEO under the accession number GSE168298 and are publicly available as of the date of publication.

## ACKNOWLEDGMENTS

We acknowledge ZVM team for their expert animal care. This work was supported by RUB/FoRUM (F1008N-20, IF-027N-22) grants; German Research Foundation (DFG) grants TU432/3-1 and TU432/6-1, Schram-Stiftung (TT); GRK2862/1/492434978 (HPN); DFG grant through the Heisenberg Program (AR 732/3-1) and Germany’s Excellence Strategy (CIBSS – EXC-2189 – Project ID 390939984) (SJH). The authors declare no competing financial interests.

## AUTHOR CONTRIBUTIONS

GS, PAU, HDN, LP, VTC and PK analyzed RNA-seq data, generation, and validation of sgLncSox1 and mouse phenotype. HPN, SJA and BBS provided research tools and offered discussions for the study; TT conceived the study; TT and GS wrote the manuscript.

## COMPETING INTERESTS

The authors declare no conflict of interest.

---

**Figure S1.**
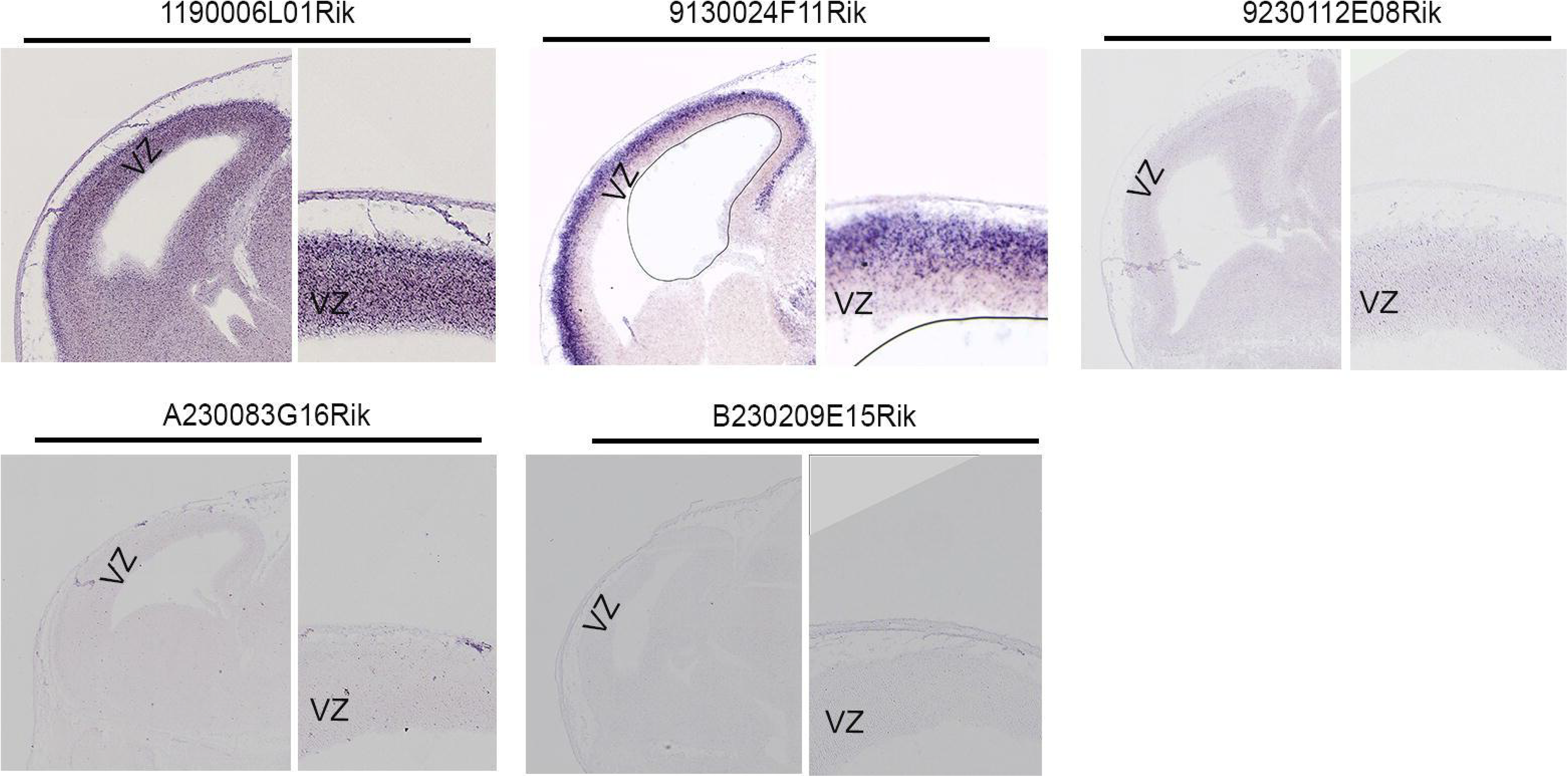

**Figure S2.**
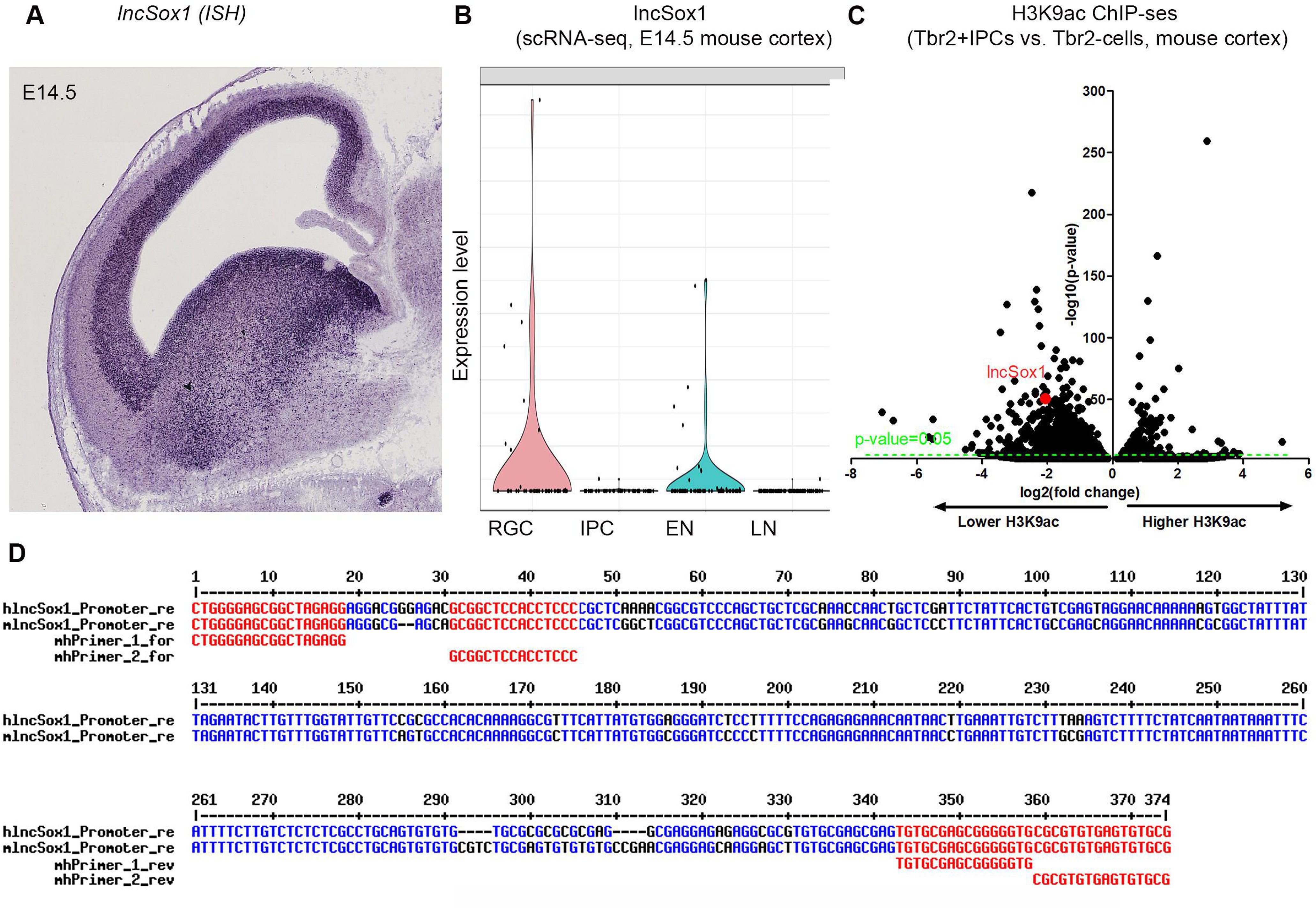

**Figure S3.**
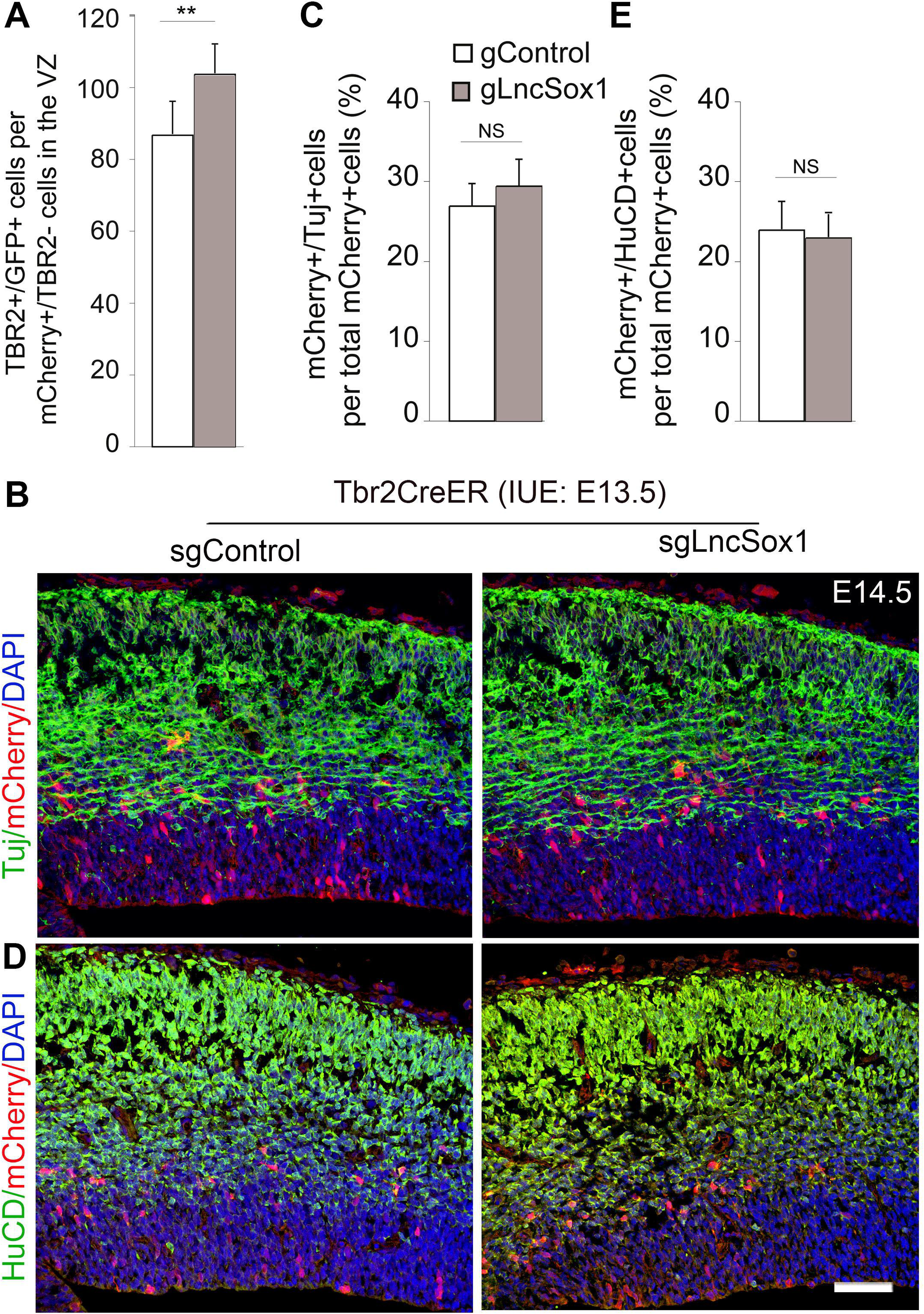

## Notes

### Competing Interest Statement

The authors have declared no competing interest.

