## Supplemental information for "Targeted H3K9 acetylation at lncSox1 promoter by cell type-specific epigenome editing promotes intermediate progenitor proliferation in developing mouse cortex"

***Supplemental figures***

**Figure S1: Expressions of ncRNAs within and outside the ventricular zone of the developing mouse cortex.**

Micrographs obtained from GenePaint showing *in situ* hybridization in the developing mouse cortex and indicating that whereas some ncRNAs are expressed in the ventricular zone, others are preferentially expressed in the (presumptive) cortical plate.

**Figure S2: Expression of lncSox1 in mouse cortex is mainly found in non-IPCs and associated with low promoter levels of H3K9ac.**

(A) Micrograph showing *in situ* hybridization (ISH) of *lncSox1* in the E14.5 mouse cortex with overt signal intensity in the ventricular zone. (B) Image showing single cell RNA sequencing analysis of *lncSox1* expression profile in the E14.5 mouse cortical cells. (C) Volcano plots showing H3K9ac distribution profile of IPC and non-IPC genes following ChIP/qPCR sequencing analysis. The *lncSox1* gene is highlighted in red. (D) DNA sequence alignment for two ortholog regions of mouse (m) and human (h) *lncSox1* promoter and primers, which were used in ChIP/qPCR experiment (see also Fig. 3E). Note that mhPrimer_1 set was used to amplify the region 1 of both mouse and human *lncSox1* promoter. The mPrimer_2 and hPrimer_2 sets, which have similar sequence, were used to amplify the region 2. Abbreviations: RGC, radial glial cell; IPC, intermediate progenitor cell; EN, early-born neuron; LN, late-born neuron.

**Figure S3: H3K9ac-mediated increase in lncSox1 expression promotes generation of IPCs without altering early-stage neurogenesis**

(A, B) Bar graph (A) and immunomicrograhs (B) showing significant increase in Tbr2 expressing cells following treatment of the E14.5 mouse cortex with sgLncSox1 compared to control. (C-E) Bar graphs (C, E) and immunomicrograhs (D) showing no significant difference in the number of early-born neurons following treatment of the E14.5 mouse cortex with sgLncSox1 compared to control. Values are presented as means ± SEMs (***p* < 0.001, ns: not significant). Scale bar = 50 µm.

***Supplemental data files***

- Table S1: Differential binding of H3K9ac at ncRNA promoters between TBR2+ IPCs in TSA- and vehicle-treated embryonic cortex (as a Supplemental Spreadsheet).
- Table S2: Differential ncRNA gene expression between TBR2+ IPCs in TSA- and vehicle-treated embryonic cortex (as a Supplemental Spreadsheet).
- Table S3: Differential binding of H3K9ac at ncRNA promoters between TBR2- cells in TSA- and vehicle-treated embryonic cortex (as a Supplemental Spreadsheet).
- Table S4: Differential ncRNA gene expression between TBR2- cells in TSA- and vehicle-treated embryonic cortex (as a Supplemental Spreadsheet).
- Table S5: statistical analyses (as a Supplemental Spreadsheet).
- Table S6: List of primers for qPCR and ChIP-qPCR (as a Supplemental Spreadsheet).
